# Aberrant brain-type neuronal programs in large-cell pancreatic neuroendocrine carcinoma

**DOI:** 10.1101/2025.06.18.660360

**Authors:** Olivia Debnath, Hilmar Berger, Katharina Detjen, Philipp Kirchner, Ilaria Mariononi, Ines Eichhorn, Felix Bolduan, Thomas Conrad, David Horst, Roland Eils, Bertram Wiedenmann, Aurel Perren, Naveed Ishaque

## Abstract

High-grade pancreatic neuroendocrine carcinoma (panNEC) is a rare, aggressive cancer with limited tissue availability. Single-nucleus transcriptomics of five large-cell panNECs revealed two clinically relevant neuroendocrine cell states: one highly proliferative with an aberrant brain-specific neuronal differentiation program and another stress-responsive state enriched for heat stress, hypoxia, and glycolysis. Our findings suggest potentially new therapeutic vulnerabilities, highlighting the need to evaluate the efficacy of combination therapies.

## Main

Pancreatic neuroendocrine carcinoma (panNEC) is a rare, highly aggressive tumor defined by a poorly differentiated neuroendocrine morphology, expression of neuroendocrine markers, and Ki67 index >20%, clinically very different from well-differentiated pancreatic neuroendocrine tumors (panNET) [1]. Despite aggressive treatment, median overall survival for metastatic gastroenteropancreatic NECs is <1 year [2]. panNEC manifests as large-cell (LC-panNEC) or small-cell (SC-panNEC) [1], with LC-panNECs displaying greater heterogeneity and less predictable response to platinum-based chemotherapy [2]. Molecular characterization of LC-panNECs remains challenging due to limited tumor tissue access and the lack of experimental models. Frequent systemic spread often precludes surgery, further restricting tissue availability. Bulk omics profiling suggests panNECs share features with normal and neoplastic pancreatic exocrine lineages [3, 4] and exhibit genomic and epigenomic similarities to acinar or ductal cells [3, 5], complicating their biological classification. To resolve this heterogeneity, we performed single-nucleus transcriptomics on five resected LC-panNEC cases using fresh-frozen tissue, generating a detailed cellular map to uncover malignant cell states and potential therapeutic targets.

We analyzed tissues of five patients with LC-panNEC, including one metastatic sample (P1) pre-treated with Cisplatin/Etoposide and FOLFIRINOX **(Figure 1a & Extended Data Figure 1).** We isolated and sequenced single nuclei from fresh frozen specimens, resulting in 45,015 high-quality nuclei after quality control **(Supplementary Table 1).** Data integration and clustering revealed that nuclei from four patients (P1-P4) clustered together, while those from patient P5 segregated, indicating a distinct transcriptomic profile and cellular composition **(Figure 1b, Extended Data Figure 1).** Using cluster-specific marker genes, we defined five main cell types: neuroendocrine (NE), amphicrine acinar, amphicrine progenitor-like, normal stroma, and immune cell types **(Figure 1c)**. Both NE and amphicrine cells expressed canonical NE diagnostic markers *CHGA, SYP,* and *NCAM1* (**Figure 1d**). Immunohistochemical validation confirmed high protein expression of CHGA and SYP across all five samples, while INSM1 expression, a transcription factor essential for neuroendocrine differentiation [6], was detected only in P1–P4 but not P5 **(Extended Data Figure 1b).** Cells from the NE cell type were detected across all patient samples (P1–P5), predominating in P1–P4 while present at a lower frequency in P5 **(Extended Data Figure 1e, f)**. The P5-associated amphicrine acinar and progenitor-like cell types expressed mature acinar (*GP2, RBPJL, MECOM, NR5A2, CEL, PRSS1, PRSS2*) or pancreatic progenitor features (*PDX1*), respectively. Stromal cells expressed the mesenchymal marker *VIM* (**Figure 1d**) and extracellular matrix (ECM) remodeling genes **(Extended Data Figure 1g).**

**Figure 1.**
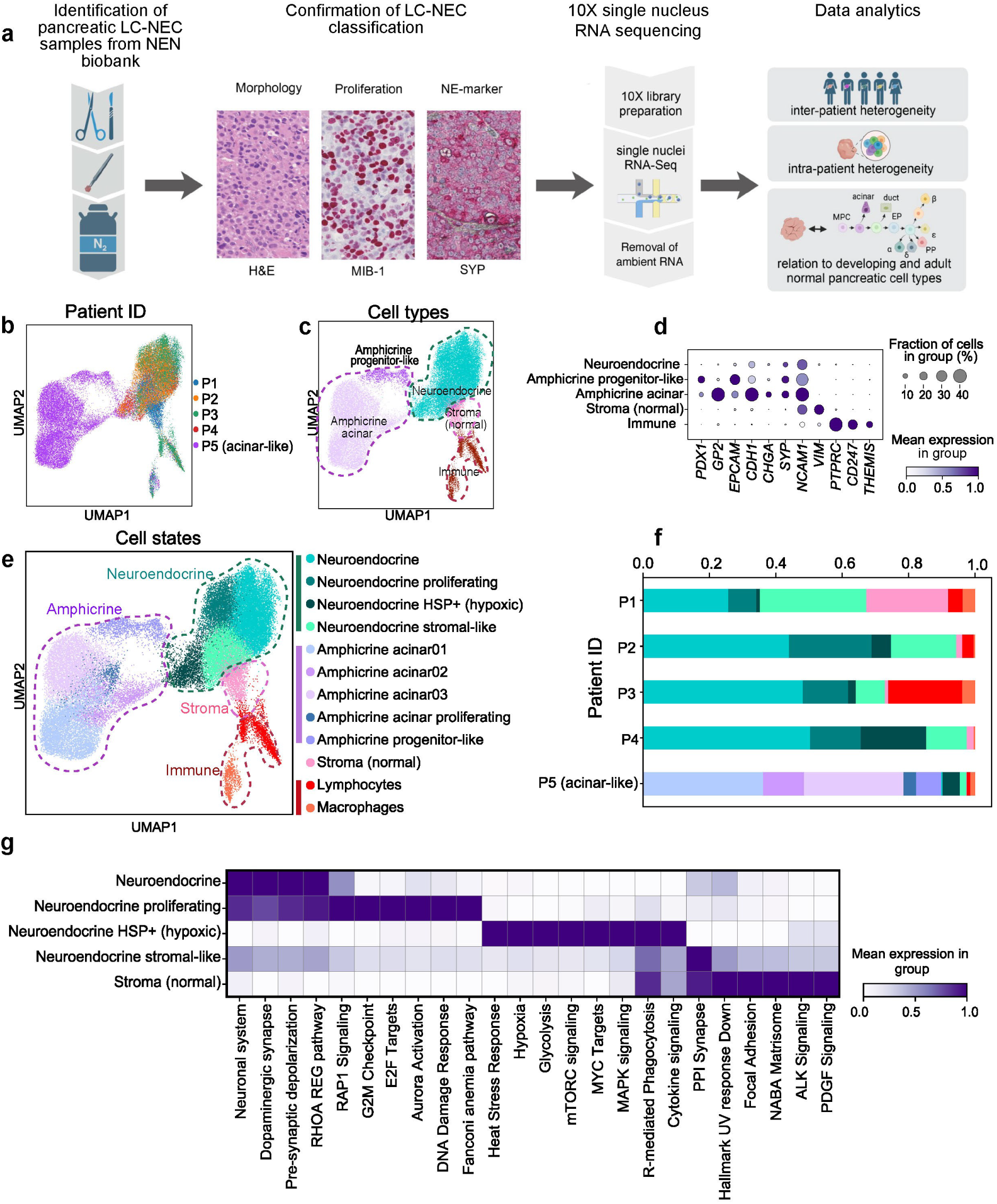
panNEC cellular landscape reveals unique and shared cell states. (a) Study design schematic. UMAP of cellular communities, color-coded by (b) Patient ID and (c) cell types from snRNA-seq. (d) Dot plot of neuroendocrine and lineage markers. (e) UMAP of cell states. (f) Stacked bar plot showing relative composition of inferred cell states per patient. Unique cell-states are specific to P5. Color legend shared with (e). (g) Heatmap of differentially expressed pathways in shared cell states (refer to Supplementary Table 3).

**Extended Data Figure 1.**
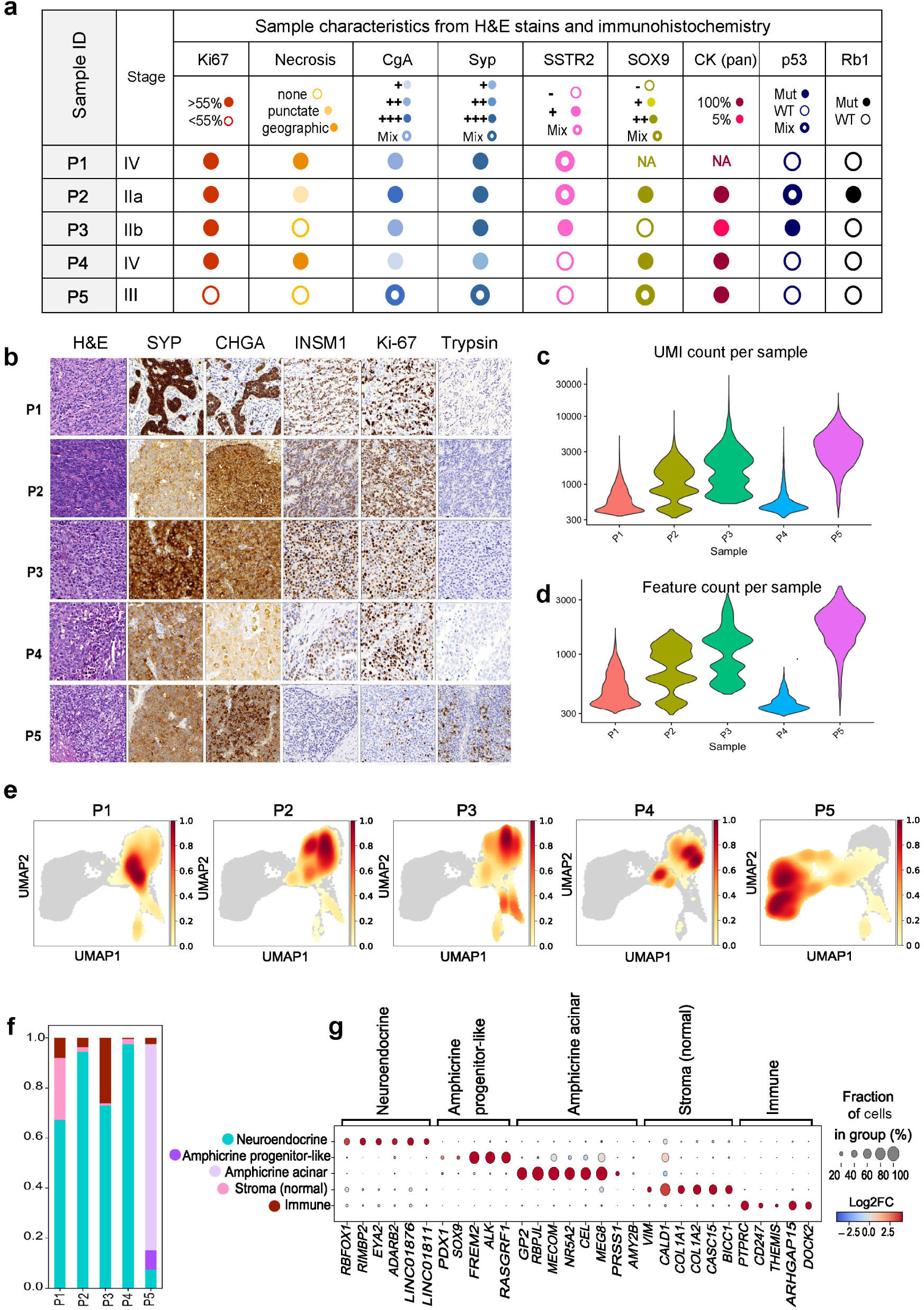
Sample characterization and quality control of panNEC samples. (a) Table summarizing sample characteristics, including tumor stage, Ki-67 index, necrosis status, neuroendocrine marker expression (CgA, SYP, SSTR2, SOX9, pan CK), and mutation status of p53 and Rb1. (b) Representative images of H&E, Synaptophysin, Chromogranin, INSM1, Ki-67, and trypsin immunostaining, highlighting tumor heterogeneity in sample P5, which shows amphicrine differentiation. (c) Violin plots depicting total UMI counts and (d) expressed genes per sample. (e) Gaussian Kernel density estimation of nuclei distribution in UMAP embedding across panNEC samples. (f) Stacked bar plot showing proportions of cell types per patient. (g) Dot plot of specific markers for each cell type, color-coded by log2FC and size-coded by percentage of nuclei expressing each marker. Manually added PDX1 and SOX9 markers show positive log2FC in amphicrine progenitor-like cells.

The neuroendocrine (NE) cell type present was further refined into four distinct cell states: NE, NE proliferating, NE HSP+ (hypoxic), and NE stromal-like, each mapping to specific functional pathways and oncogenic processes **(Figure 1g & Supplementary Tables 2, 3).** The relative abundance of these shared NE cell states varied considerably between individual NEC cases **(Figure 1f & Extended Data Figure 1e)**. Both the NE and NE proliferating cell states prominently expressed neuronal markers *RBFOX1, RIMBP2,* and *ADARB2*, and were enriched for voltage-gated Ca2+ and K+ channel genes (*CACNB2, CACNA1A, KCNJ3,* and *KCNJ6*), suggestive of pre-synaptic depolarization **(Extended Data Figure 2a).** *RBFOX1* is not expressed in normal embryonic or adult pancreas [7], suggesting aberrant expression in NEC. RBFOX1 promotes neuroplasticity [8] and regulates calcium signaling [9]. The NE proliferating cell state expressed *MKI67, DIAPH3*, and *CENPP* and enrichment of cell-cycle, proliferation, and DNA damage response (DDR) pathways, including Fanconi anemia (FA) signaling pathway, regulated by *FANCA, FANCI*, tumor suppressor genes *BRCA1/2,* and the *BRCA-*interacting protein-encoding gene BRIP1 **(Figure 1g & Extended Data Figure 2a).** In contrast, the NE HSP+ (hypoxic) cell state expressed high levels of heat-shock protein (HSP) genes (*HSP90AA1, HSP90AB1, HSPE1 & HSPH1*) alongside hypoxia-induced genes (*VEGFA, KDM3A, NDRG1*) **(Extended Data Figure 2a)**. This cluster was enriched in pathways related to heat stress response, hypoxia, mTORC signaling, glycolysis, upregulated MYC targets, and MAPK signaling (**Figure 1g**). Further details on the remaining cell states are provided in the Supplementary Information.

Gene set enrichment analysis (GSEA) revealed therapeutic vulnerabilities amenable to approved or investigational drugs (**Extended Data Figure 2b-g**). Cisplatin emerged as a key inhibitor of the NE proliferating cell state, suggesting that first-line treatments effectively target this state while sparing others. In addition, NE proliferating cells exhibited elevated *EZH2* expression (**Extended Data Figure 2a, h**), a marker linked to advanced pancreatic neuroendocrine neoplasm (panNEN) progression [10]. However, a small *EZH1/2* co-expressing subpopulation within NE proliferating cells (**Extended Data Figure 2h, i**) might exhibit resistance to EZH2 inhibition through functional redundancy of the PRC2 complex. In light of this heterogeneity, dual EZH1/2 targeting (e.g., Valemetostat) could offer a more effective therapeutic strategy than EZH2 inhibitors alone.

In contrast, the NE HSP+ (hypoxic) cell state appeared susceptible to HSP90 inhibitors, including Geldanamycin and Tanespimycin. Moreover, the NE HSP+ (hypoxic) state exhibited high *HSF1, HIF1A*, and *ATF5* transcriptional activity (**Extended Data Figure 3a**), reflecting a stress-adapted, non-neuronal state. HSF1 is known to promote cancer growth and is associated with poor prognosis, while its co-regulation with HIF1A provides a mechanistic link between hypoxia and heat-shock activation [11]. *ATF5,* essential for cancer survival and proliferation, is associated with poor outcomes across malignancies [12].

**Extended Data Figure 2.**
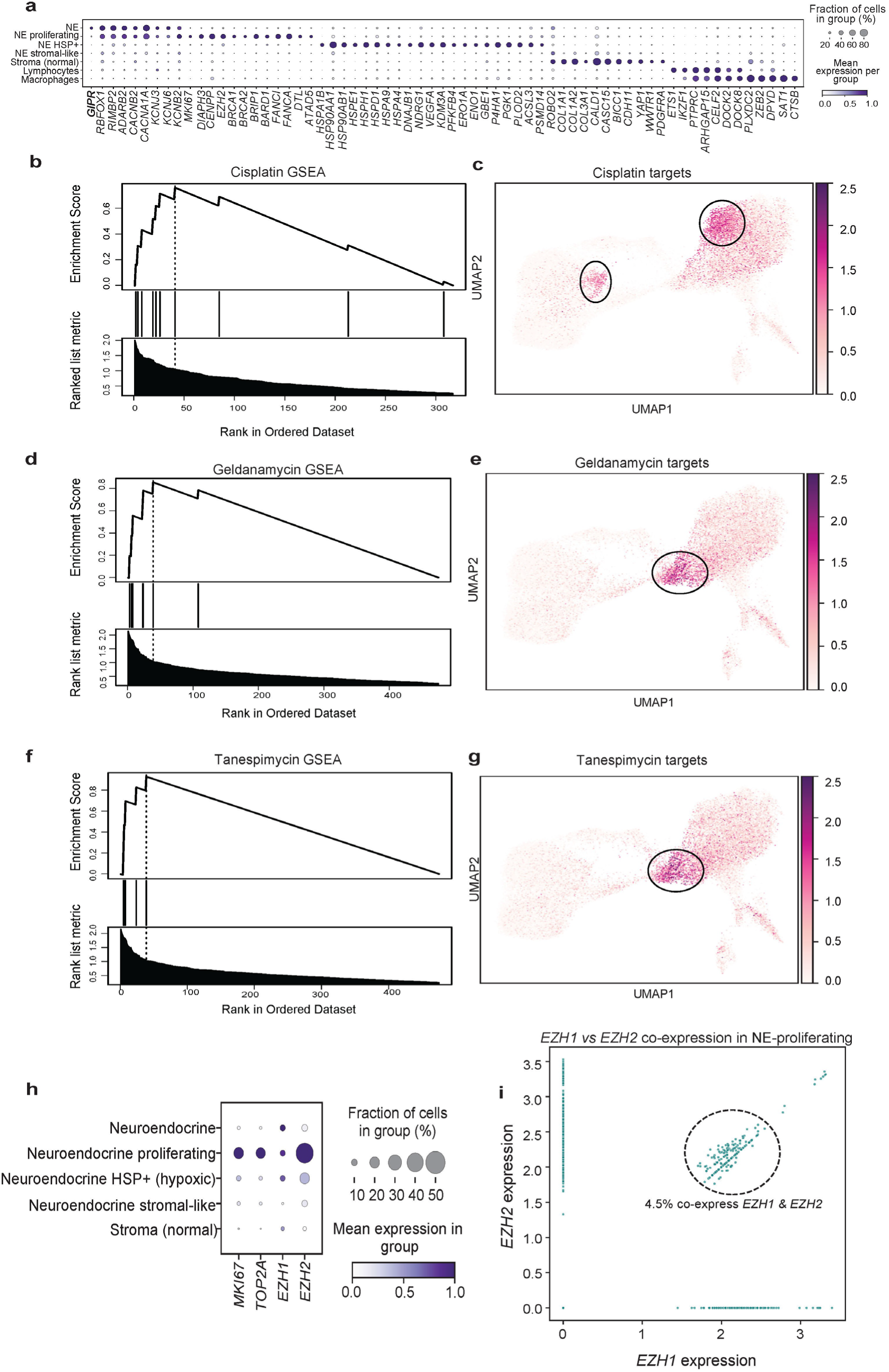
Shared NE proliferating and NE HSP+ (hypoxic) cell states are pathological hubs with therapeutic vulnerabilities. (a) Dot-plot illustrating selected markers across shared panNEC cell states (average log2FC ≥ 0.25, Bonferroni adjusted p-value < 0.01; **Supplementary Table 2**). Gene expression is color-coded, and marker size reflects the percentage of nuclei expressing each gene. Expression levels are standardized for comparison. Cells most enriched for druggable pathways are circled for convenience. GIPR (shared NE) was manually included as it represents a potential therapeutic target in P3; YAP1/WWTR1/PDGFRA were inferred using LR, as well as represent therapeutic vulnerabilities in P1. NE= Neuroendocrine. NE HSP+ (hypoxic) is mentioned as NE HSP+. GSEA identified drug targets: (b) Cisplatin for NE proliferating (NES= 2.0493 & FDR= 0.005223), (d) Geldanamycin (NES= 2.1501 & FDR= 0.0043136), and (f) Tanespimycin (NES= 2.0704 & FDR= 0.003505) for NE HSP+ (hypoxic). Feature plots display nuclei enriched for (c) Cisplatin, (e) Geldanamycin, and (g) Tanespimycin targets.

We then investigated the developmental origins of LC-panNEC. Lack of expression of pancreatic endocrine lineage markers (*GCG, INS,* and *SST)* confirms that panNEC cell states do not recapitulate mature islet profiles (**Extended Data Figure 3b; Supplementary Table 4**). They also showed no enrichment for acinar or ductal markers, suggesting an absence of key exocrine pancreatic features. Regulon activity analysis in NE and NE-proliferating states identified transcription factors linked to pancreatic and brain organogenesis, including *PAX6, ISL1, SOX5, ETV1,* and *TCF3* (**Figure 2a**). *PAX6* regulates islet and neuronal development [13–15], while *ISL1* governs islet development, motor neuron generation, and cranial ganglia formation [16]. Their activities were elevated in panNEC cell states compared to their inferred physiological levels in the pancreas, reinforcing their transcriptional divergence from normal pancreatic lineages (**Figure 2a**). Moreover, GTEx-based differential gene expression analysis identified mutually exclusive gene signatures, with amphicrine states enriched for pancreatic markers, while shared NE states preferentially expressed brain-associated programs (**Figure 2b**).

**Figure 2.**
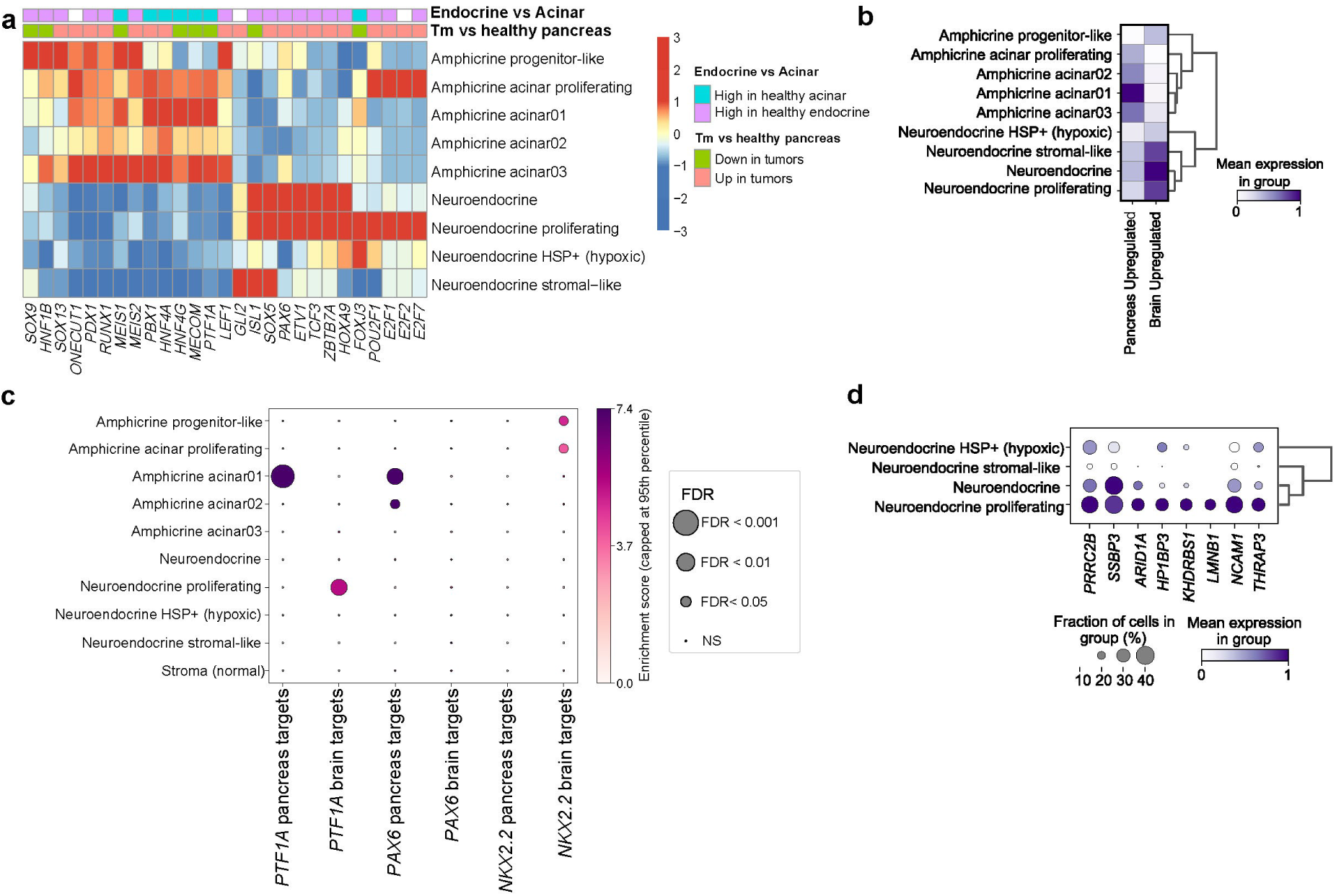
PTF1A-directed brain targets characterize aberrant neuronal phenotype in the neuroendocrine proliferating cell state. (a) Heatmap depicting differentially regulated transcription factor activities across amphicrine and shared panNEC cell states predicted by pySCENIC (Z-score expression of targets). (b) Matrix-plot showing pancreas and brain signature score across panNEC cell states inferred using DESeq2 analysis and GTEX (see Methods). (c) Dot plot showing enrichment of PTF1A, PAX6, and NKX2.2 target genes derived from pancreas and brain. Enrichment scores (representation factor, capped at the 95th percentile for visualization) are color-coded; FDR is size-coded. Stroma (normal) is included as a negative control to illustrate that enrichment is specific to epithelial-derived tumor cells, and not broadly observed across the tumor microenvironment (TME). Rationale for using PAX6 and NKX2.2 is explained in the Supplementary Information. (d) Dot plot showing differentially upregulated markers contributing to PTF1A brain signature over-enrichment in the NE proliferating cell state. Average gene expression is color-coded, and the percentage of nuclei expressing each marker is size-coded. Stroma (normal), amphicrine, and immune cell states are excluded.

NE and NE-proliferating markers showed significant enrichment within brain-specific DEGs (comparing brain vs. normal pancreas, see Methods), implicating *REST, GFI1,* and *TEAD4* as major transcriptional regulators (**Extended Data Figure 3c, d**). *REST*, a key repressor of neuronal genes, governs neuroendocrine differentiation in cancer [17]. Next, human- developing brain atlas data [18] was mapped onto the panNEC data set, and the shared NE cell states were annotated as subcortical neurons (**Extended Data Figure 3e**). These cell states are marked by *RBFOX1*, which regulates synaptic connectivity, axon guidance, and neuronal excitability, and is not expressed in adult pancreas cell types [19] (**Extended Data Figure 3f, g**). Orthogonally, *RBFOX1* showed remarkably robust expression in GABAergic cortical interneurons and L2/3-6 intratelencephalic projecting glutamatergic neurons across large-scale brain single-cell datasets [20] (n ≈ 1.13 million and 1.79 million single cells/nuclei, respectively; **Supplementary Table 5**), reinforcing the ’brain-specific’ identity of these NE cell states.

Previous investigations of panNEC using DNA methylation and mutational profiles indicate an exocrine cell origin and a mutational spectrum similar to exocrine PDAC rather than well-differentiated NET [3, 5]. We observed significant *PTF1A* regulatory activity in the amphicrine acinar cell states (**Figure 2a**), consistent with its established role in exocrine pancreatic development and acinar lineage specification [21–25]. Notably, *PTF1A* is also a critical regulator of brain development [26–29]. Building on previous findings that linked shared panNEC cell states to brain-like signatures, we explored the dual relevance of *PTF1A* in regulating pancreas- and brain-specific transcriptional programs. Using murine tissue-specific developmental signatures [28], we found that *PTF1A*-regulated brain targets were significantly enriched in the NE proliferating cell state, including genes such as *HP1BP3, KHDRBS1, SSBP3, THRAP3, PRRC2B, ARID1A, LMNB1,* and *NCAM1*, while *PTF1A*-regulated pancreatic targets were enriched exclusively in the amphicrine acinar 01 cell state (**Figure 2c**, **Supplementary Table 4**). Apart from *NCAM1*, none of the *PTF1A* brain targets were expressed in adult pancreas cell types or states, suggesting their role in neuronal dedifferentiation and maintenance (**Extended Data Figure 3h**). Notably, NE proliferating cells expressed *HP1BP3* and *EZH2*, known to drive temozolomide resistance in glioblastoma [30], suggesting a potential parallel resistance mechanism in panNEC. Additionally, *KHDRBS1*, another *PTF1A*-regulated brain target, contributes to cancer stem cell modulation, proliferation, and survival in human cancers (Figure 2d) [31, 32].

**Extended Data Figure 3.**
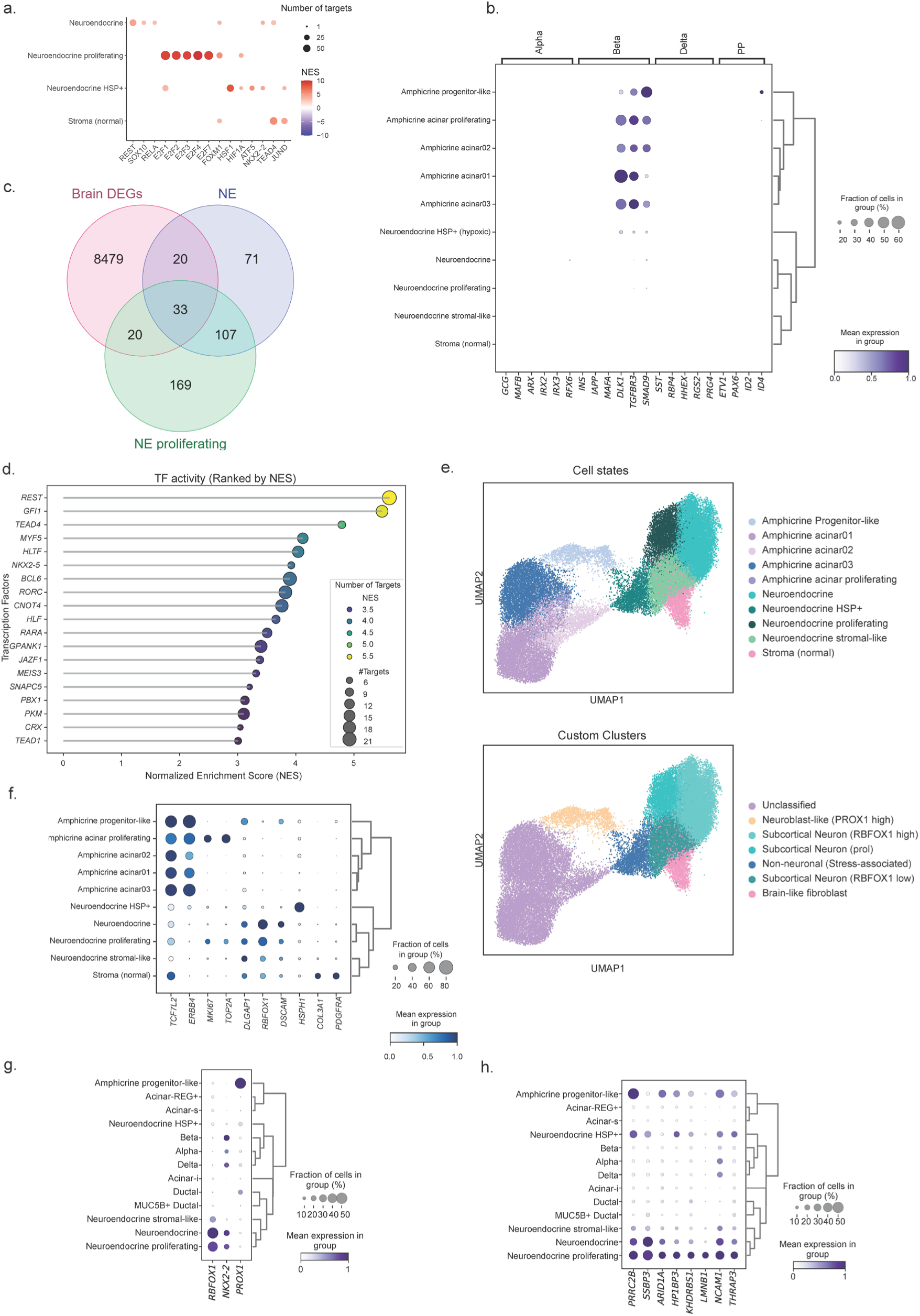
Shared NE and NE proliferating panNEC cell states recapitulate brain-specific programs. (a) Transcriptional activities inferred using iRegulon based on the top-100 differentially expressed targets (log2FC ≥ 0.25, Bonferroni adjusted p-value < 0.01). The size of each bubble reflects the number of targets regulated by a transcription factor, while the color represents normalized enrichment scores (NES). (b) panNEC cell states lack essential endocrine lineage markers — GCG, INS, and SST. Markers were curated from Muraro et al. 2016 [34] (see Supplementary Table 4) and validated by literature curation. GCG, INS, and SST are well-established markers for alpha, beta, and gamma cell types, respectively. PPY, the characteristic marker for PP cells, was not detected in the dataset. (c) Venn diagram illustrating the overlap between GTEX brain-specific (inferred by comparing brain vs pancreatic samples using DESeq2 analysis; see Methods), NE, and NE proliferating markers. The diagram displays the overlap between differentially expressed genes from brain vs pancreas tissue (see Methods), NE markers, and NE proliferating markers (log2FC ≥ 0.25, Bonferroni adjusted p-value < 0.01; Logistic Regression). (d). Lollipop plot depicting transcription factor (TF) activity inferred from shared neuroendocrine (NE) and NE-proliferating markers overlapping with brain-specific differentially expressed genes (DEGs) identified using DESeq2 analysis. TF activity was predicted using iRegulon, and TFs are ranked by their normalized enrichment scores (NES) on the x-axis. Each lollipop represents a TF, with the length of the line indicating the NES. Dot size corresponds to the number of target genes associated with each TF, and dot color reflects the NES, ranging from dark blue (low NES) to yellow (high NES). REST shows the highest NES values, indicating its potential regulatory roles within the shared NE and NE-proliferating gene signatures. (e). UMAP projection of panNEC samples, where each color encodes a cell state (excluding immune cell states) and is classified into developing brain-like phenotypes. Our classification represents a semi-supervised annotation of panNEC cell states using a developing brain atlas reference [35]. Most shared NE cell states are classified as subcortical neurons (where NE & NE proliferating are RBFOX1 high). (f). Dot plot showing key selected genes across classified cell states, showing drivers of brain-like phenotype (see **Methods**). Gene expression is color-coded, and marker size reflects the percentage of nuclei expressing each gene. (g) Dot-plot depicting normalized expression of RBFOX1, NKX2-2, and PROX1 across panNEC cell states (purple text) and the adult pancreas dataset from Tosti et al. 2021 [19]. Average gene expression is color-coded, and size indicates the percentage of nuclei expressing each gene (capped at 40%). RBFOX1 is upregulated in NE and NE proliferating cell states, while PROX1 is significantly elevated in the amphicrine progenitor-like sub-state **(Supplementary Table 2).** (h) Dot plot highlighting key PTF1A-regulated brain targets across panNEC cell states and adult pancreas cell types/states, integrated from Tosti et al. 2021 [19]. Adult islet cells (alpha, beta, and delta) show minimal expression of PTF1A-regulated brain targets. P5 amphicrine acinar cell states were excluded from the plot. Gene expression is color-coded based on average expression within each cluster, and marker size represents the percentage of nuclei expressing each gene. Gene expression levels are standardized across columns for comparison. Hierarchical clustering of all expressed genes in the concatenated dataset illustrates global transcriptomic similarities.

**Extended Data Figure 4.**
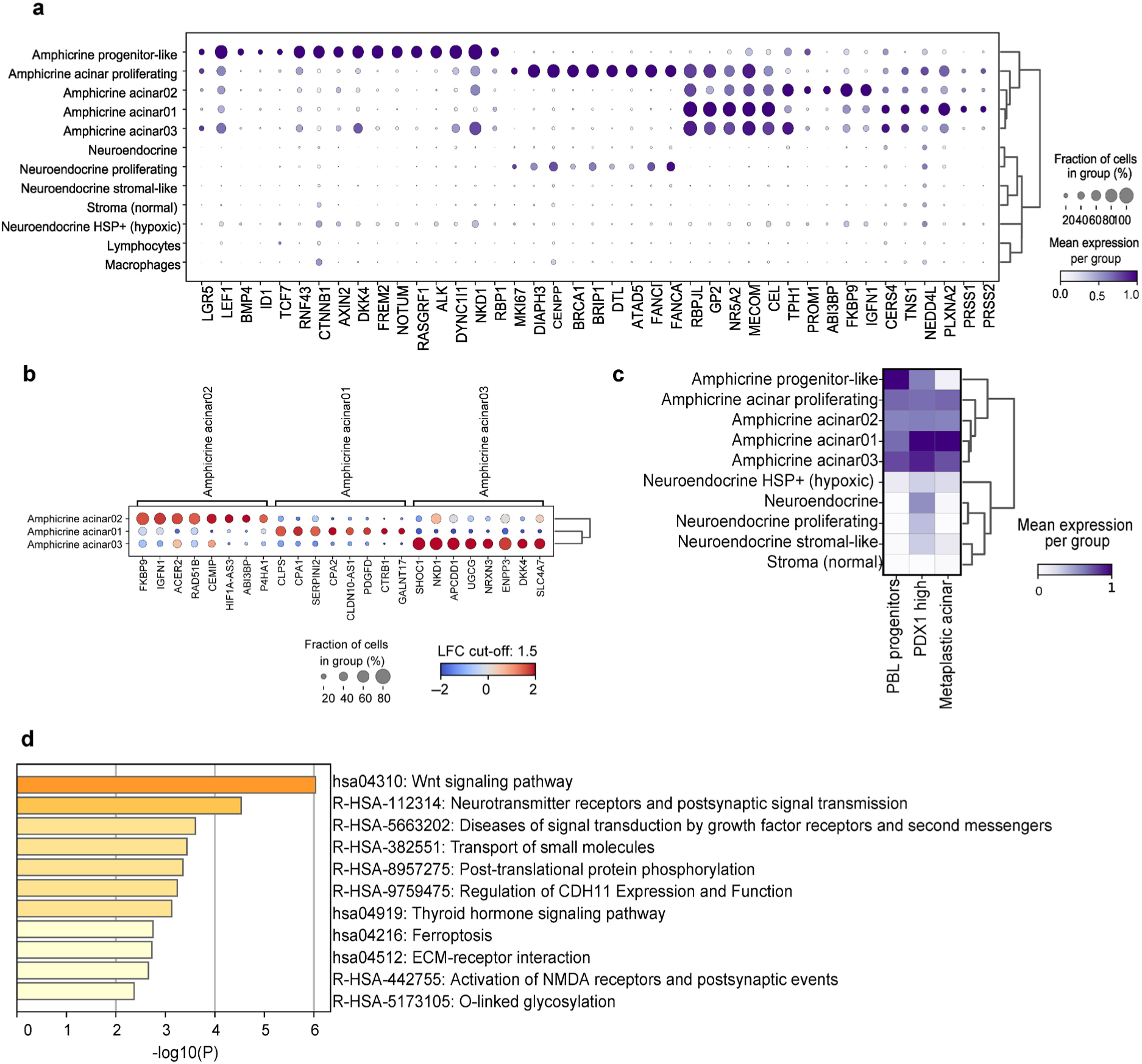
Amphicrine progenitor-like sub-state is characterized by WNT-BMP-NOTCH signaling. (a) Dot-plot showing key markers for amphicrine progenitor-like and acinar cell states (log2FC ≥ 0.25, Bonferroni p < 0.01). Only markers specific to amphicrine cell states are displayed. Average gene expression is color-coded, and marker size reflects the percentage of nuclei expressing each gene. (b) Dot plot showing differentially expressed markers across amphicrine acinar cell states. DEGs were recalculated specifically for amphicrine acinar cell states using the Wilcoxon rank sum test (avg log2FC ≥ 1.5, Benjamini-Hochberg corrected p-value < 0.01). Dot color represents the average log2FC per marker within each sub-state, while the dot size indicates the percentage of cells expressing the marker. (c) Matrix plot of module scores for PBL progenitors (PDX1 high pancreatic cells) and metaplastic acinar cells. (d) Bar graph of enriched pathways from upregulated markers of the amphicrine progenitor-like sub-state.

In conclusion, our study reveals that shared neuroendocrine cell states in LC-panNEC exhibit brain-type rather than islet differentiation, consistent with existing mutational and epigenetic evidence [3, 5]. Notably, we identified brain-associated programs in NE and NE-proliferating cell states, resembling subcortical neurons with excitatory properties. *PTF1A*-regulated brain-specific transcriptional signatures within the proliferative state suggest that LC-panNEC co-opt neuronal transcriptional networks to enhance aggressiveness, stemness, and therapy resistance, unveiling novel therapeutic vulnerabilities. In addition, the hypoxia-adapted NE HSP+ (hypoxic) cell state exhibited heat stress responses, hypoxia, and metabolic reprogramming. This state likely reflects cancer cells’ reliance on HSP90 for stabilizing oncoproteins, a phenomenon known as “HSP90 addiction” [33], and mirrors conserved tumor stress responses across cancers.

Collectively, our findings pinpoint shared therapeutic vulnerabilities, where combining HSP90 inhibitors with Cisplatin or EZH2 inhibitors could jointly target neuroendocrine HSP+ (hypoxic) and proliferative cell states. These strategies warrant rigorous validation and may shift therapy from molecular marker targeting to disrupting malignant cell states, advancing precision medicine, and improving LC-panNEC outcomes.

## Methods

### Patient cohort and ethics approval

Due to the rarity of the disease, a retrospective study design was utilized to enable strict sample inclusion criteria. Inclusion criteria required histopathological confirmation of G3 NEC with large cell morphology, tumor localization in the pancreas, and availability of both fresh frozen and formalin-fixed, paraffin-embedded (FFPE) tumor tissue. A cohort of six patients diagnosed with high-grade GEP-NEN was selected from two ENETS Centers of Excellence: the University Hospital Charité Berlin, Germany, and the University Cancer Institute of the Inselspital and the University of Bern, Switzerland. Samples were reviewed and (for older samples) reclassified according to WHO 2019 criteria by a board-certified pathologist (Prof. Aurel Perren), as detailed in Table 4. TNM staging was provided based on the 8th edition of the Union for International Cancer Control (UICC)/American Joint Committee on Cancer (AJCC) guidelines. Ethical approval was obtained from local authorities (Ethics approval EA1/229/17 at Charité and KEK BE 105/2015 from Kantonale Ethikkommission Bern), in accordance with the Declaration of Helsinki. Patient samples were anonymized for analysis. The mutational status of samples was determined by prior panel sequencing by April-Monn et al. [4].

Both FFPE and fresh frozen tumor tissues were obtained. FFPE sections were stained with H&E and subjected to immunohistochemical detection of chromogranin A, synaptophysin, insulinoma-associated protein 1, pan-cytokeratin, trypsin, Ki67, TP53, Rb1, and Sox9, using standard clinical diagnostic protocols available at the Institute of Pathology, Bern University. Single-nucleus RNA-seq (snRNA-seq) sample preparation was conducted on fresh frozen tissues at three separate time points: the first run used P1 as a pilot for technical feasibility, the second run included samples P2-P6 (however, sample P6 failed QC and was excluded from subsequent analyses), and the third run repeated the P5 preparation due to poor quality observed initially, processed in two batches (technical replicates).

### panNEC Immunohistochemistry (IHC) validation

All IHC markers were repeated on freshly cut tissue blocks and re-evaluated by a neuroendocrine neoplasms (NEN) expert pathologist (Prof. Aurel Perren). Paraffin-embedded tissue was sectioned into 2.5 µm-thick serial sections, then deparaffinized, rehydrated, and subjected to antigen retrieval using an automated immunostainer (Bond RX, Leica Biosystems). Antigen retrieval was carried out in either Tris-EDTA buffer or Citric acid buffer, typically for 30 minutes at 95°C, depending on the specific marker (see table below for antibody details). Primary antibody incubation was performed for 30 minutes at the specified dilutions. Visualization was achieved using a Bond Polymer Refine Detection Kit (Leica, #DS9800) (RRID: AB_2891238), with DAB (3,3′-Diaminobenzidine) as the chromogen. Slides were counterstained with hematoxylin. Scanning was conducted using an automated slide scanner (Panoramic 250, 3DHistech) at 20x magnification, and images were captured using QuPath software.

| Antibody | Supplier | ID | Dilution | Antigen Retrieval | Temperature | Time |
| --- | --- | --- | --- | --- | --- | --- |
| CgA | CellMarque | 138M-94<br>LK2H10 | 1:400 | Tris-EDTA | 100°C | 30 min |
| Ki-67 | Dako | M7240 | 1:200 | Tris-EDTA | 95°C | 30 min |
| RB1 | BD Pharmingen | 554136 G3-245 | 1:200 | Citric buffer | 100°C | 30 min |
| SOX9 | Cell Signaling | 82630 T<br>D8G8H | 1:100 | Tris-EDTA | 95°C | 30 min |
| SSTR2A | BioTrend | SS-8000-RM UMB-1 | 1:50 | Tris-EDTA | 100°C | 30 min |
| Synaptophysin | Novocastra | 27G12 | 1:100 | Tris-EDTA | 100°C | 30 min |
| TP53 | Dako | M7001 DO-7 | 1:800 | Citric buffer | 95°C | 20 min |
| Trypsin 1 | Chemicon | MAB1482 | 1:20000 | Proteinase K solution | 37°C | 5 min |
| EZH2 | Cell Signaling | 5246 | 1:400 | Citric buffer | 100°C | 30 min |
| INSM1 | Santa Cruz | Sc-271408 | 1:200 | Tris-EDTA | 95°C | 40 min |
| Pan-Cytokeratin | DAKO | M3515 | 1:400 | Tris-EDTA | 95°C | 30 min |
| YAP1 | Abcam | Ab52771 | 1:200 | Tris-EDTA | 95°C | 30 min |

### panNEC sample preparation and snRNA-seq experiment

Tumor tissue preserved by cryopreservation was disrupted in NP-40 lysis buffer (10 mM Tris-HCl at pH 7.4, 10 mM NaCl, 3 mM MgCl₂, 0.01% NP-40, 1 mM DTT, 2% BSA, 1 U/µl RNAse inhibitor, and Complete EDTA-free protease inhibitor) in a 1.5 ml Eppendorf tube using a plastic pestle. After a 5-minute incubation on ice, the suspension was filtered through a 70 µm pre-separation strainer, then centrifuged at 4°C and resuspended in nuclei wash buffer (PBS, 1% BSA, and 0.4 U/µl RNAse inhibitor). DAPI (3.5 µl) was added, and the mixture was passed through a 40 µm Flowmi cell strainer prior to sorting on a BD Aria Fusion instrument into an Eppendorf tube containing 200 µl of sort buffer (PBS, 2% BSA, 1 U/µl RNAse inhibitor) using a 100 µm nozzle.

Single-nuclei libraries were generated according to the protocol outlined in the Chromium Next GEM Single Cell 3ʹ Reagent Kits v3.1 (Dual Index) user guide (CG0003154) from 10x Genomics. In brief, individual nuclei were encapsulated with gel beads containing barcoded primers in the Chromium controller. Within each droplet, mRNA was reverse-transcribed into barcoded cDNA, which was then amplified. The amplified cDNA was subjected to fragmentation, end-repair, A-tailing, adaptor ligation, and sample index PCR. The resulting library was sequenced on a NovaSeq 6000 system (Illumina) to a depth of 40,000 mean reads per nucleus.

### panNEC snRNA-seq data analysis

#### Data pre-processing and Quality Control analysis

Single-nucleus RNA reads were aligned to the GRCh38-2020-A reference using CellRanger v6.0.1 [36], incorporating both exonic and intronic (nuclear pre-mRNA) reads. One biological sample was excluded from downstream analysis due to failure to meet basic CellRanger QC criteria. Sample P5 was processed as two technical replicates, which were subsequently merged during analysis. The background reads were removed using the R package SoupX [37], with parameters set to tfidfMin = 0.6, soupQuantile = 0.9, and priorRho = 0.1. After background subtraction, expression values were rounded to integers. Cells were filtered based on criteria including fewer than 450 or more than 4000 RNA features, over 5% mitochondrial read content, or genes with fewer than two reads across all samples. Mitochondrial and ribosomal genes were excluded from further analysis to minimize technical noise. Additionally, for joint analysis with the snRNA-seq data from Tosti et al. 2020 [19], all samples were downsampled to a median read count of 500 reads to validate major findings.

#### Data integration and clustering of samples

The SoupX-corrected AnnData matrices per patient were merged using scanpy [38], followed by log-normalization for downstream analysis. Highly variable genes (n=4000) were identified using sample ID as a batch key for lightweight batch correction and default parameters in scanpy’s highly_variable_genes() function. Subsequently, principal component analysis (PCA) was performed using tl.pca() with svd_solver=“arpack” in scanpy. This function calculated PCA coordinates, loadings, and variance decomposition, which were visualized using sc.pl.pca_overview() and heatmaps to display top features associated with each principal component. Calculation of the K-nearest neighbor (KNN) graph and UMAP visualization, with fine-tuning of the number of principal components and neighbors, resulted in well-separated patient samples or batches.

To achieve data harmonization and reduce batch effects across 5 panNEC samples, a sequential integration approach, known as chain integration, was applied. Initially, Harmony [39] (via scanpy external’s harmony_integrate function) was run using sample ID as the batch key. This function utilizes the Python version of Harmony, called harmonypy, to harmonize single-cell data stored within an AnnData object. Since Harmony corrects the Principal Components (PCs), it was run after Principal Component Analysis (PCA) but before computing the neighbor graph. The Harmony-adjusted PC1 vs PC2 graph indicated that P5 was distinct from the other four patients, largely due to acinar lineage-specific genes like GP2 encoded by PC1. However, this variation was deemed biological rather than a technical artifact, as P5’s immunohistochemistry confirmed trypsin positivity, indicating similarities with acinar cells. Subsequently, BBKNN [40] was used to construct a batch-corrected mutual neighbor graph, using sce.pp.bbknn() with parameters set to neighbors_within_batch=5, trim=80, and use_rep=’X_pca_harmony’ to consider Harmony-adjusted PC(s) for constructing a batch-balanced mutual neighbor graph.

The Leiden algorithm was then applied with a resolution parameter of 1 for clustering and inferring cell states. Clusters were merged when robust and specific markers were lacking. Leiden-12, containing n=540 nuclei, was excluded from further analysis due to its ambiguous transcriptomic profile. Case studies adjusting the number of PC(s) from n=10 to 50 showed structural changes in the data. The choice of PC=10 was considered suboptimal, yielding a less clear structure. When using a higher number of PCs (40 & 50), a distinct but ambiguous cluster (leiden-12, containing n=540 nuclei from P5) was separated on the UMAP plot. Improved batch mixing, especially among patient samples P1-P4, was observed with higher Average Silhouette Width (ASW) scores per cell type/state as the number of PCs increased to 40 or 50. Therefore, BBKNN’s default PC=50 was selected for the final analysis. UMAP was computed with random_state=0 (for reproducibility) and maxiter=200 (for optimized embedding layout and algorithm convergence). Initially, Leiden cluster 1, containing 8133 nuclei, was labeled as “Stromal/Mesenchymal.” Upon further sub-clustering at medium resolution (Leiden resolution=0.4), it was divided into two groups: Stromal-like NE (n=5668 nuclei) and Stromal (normal) (n=2465 nuclei).

Coarse-grained cell types were initially determined through Leiden clustering at a resolution of 0.3, and subsequently fine-tuned by consolidating cell states with distinct markers for amphicrine, neuroendocrine, stromal, and immune cells. Although all cell states specific to P5 were grouped under the label “Amphicrine,” the Amphicrine progenitor-like group was kept distinct due to its strong and specific expression of markers linked to pancreatic progenitors, cancer stemness, and signatures related to Wnt-Notch-BMP signaling. This observation was validated by re-running Logistic Regression marker analysis and Wilcoxon rank sum test, using scanpy’s tl.rank_genes_groups() with corr_method=’benjamini-hochberg.’

### Cell state marker analysis

Marker analysis of panNEC cell states was performed using a multivariate logistic regression (LR) generalized linear model, implemented in Seurat’s FindAllMarkers() function. In the LR analysis, the number of unique molecular identifiers (UMI), the number of genes, and the percentage of mitochondrial transcripts per nucleus were included as continuous covariates. Significant cell markers were identified among genes with a log2 fold change cut-off of 0.25 and expression in at least 25% of cells within each cluster. An adjusted p-value cut-off of 0.01 was applied following Bonferroni correction for multiple testing across states. Internal validation was also performed using the Negative Binomial method in Seurat.

For comparative analysis among specific cell states of amphicrine acinar groups, the Wilcoxon rank-sum method, implemented in scanpy’s tl.rank_genes_groups(), was used and verified with Logistic Regression. Statistically significant sub-state markers were identified based on a log2 fold change cut-off of 1.5 and expression in at least 25% of cells within each cluster. An adjusted p-value cut-off of 0.01 was applied using the Benjamini-Hochberg adjustment for multiple testing (the default correction method in scanpy). Applying the Bonferroni correction method did not change the marker results, which are visualized in Extended Data Figure 4.

### Transcriptional regulation analysis

Inference of transcriptional regulation was performed using the pySCENIC tool [41] to predict transcription factor activities on the downsampled data. The analysis used motifs from version 9 of the motif database (motifs v9), and binding sites were identified within the human genome version hg19, targeting sequences located 500 bp upstream as well as 5 kb and 10 kb centered around the transcription start site (TSS). Differences in transcription factor activity between groups were evaluated using the FindMarkers function from the R package Seurat. Transcription factor marker activities estimated by pySCENIC were integrated as a new assay, and a Wilcoxon rank sum test was applied to assess group differences. Specifically, we compared panNEC cell states to healthy pancreatic cell types from Tosti et al. 2020 [19]. If a transcription factor (TF) was expressed in healthy exocrine or endocrine cells, the differential analysis determined its upregulation in panNEC (Methods). Z-scores were applied to expression values for subsequent visualization.

### GTEx bulk RNA-seq analysis for differential gene expression analysis of brain and pancreas

Bulk RNA-seq count data and sample annotations were obtained from the GTEx portal (https://gtexportal.org/home/downloads/adult-gtex/bulk_tissue_expression, file version bulk-gex_v8_rna-seq_GTEx_Analysis_2017-06-05_v8_RNASeQCv1.1.9_gene_reads.gct.gz. Briefly, a DESeqDataSet object (dds) was constructed using DESeqDataSetFromMatrix() from the count data matrix (expr_mat) and sample metadata (colData). Differential expression analysis between the pancreas and brain tissues was performed with a likelihood-ratio test using the R package DESeq2, specifying fitType = “glmGamPoi“, full = ∼ SMTS, reduced = ∼ 1, and test = “LRT.” The fitType = “glmGamPoi” option in DESeq2 employed a generalized linear model (GLM) with a gamma-Poisson distribution to accommodate the overdispersion common in RNA-seq count data. Genes with an absolute value of the log2FoldChange < 500 and non-missing log2FoldChange values (NA) were cataloged as differentially expressed genes (DEGs). Additionally, a logFC cut-off of 2 and an adjusted p-value < 0.001 were used to identify mutually exclusive DEGs specific to pancreas and brain tissues.

*PTF1A* targets were sourced from Meredith et al. 2013 [28], and the utilized signature list is tabulated in Supplementary Table 4. DEG lists for each tissue were subsequently intersected with PAX6 and NKX2-2 target lists obtained from ENCODE transcription factor targets (https://maayanlab.cloud/Harmonizome/gene_set/EZH2/ENCODE+Transcription+Factor+Targets) and MSigDB (https://www.gsea-msigdb.org/gsea/msigdb/human/geneset/PAX6_TARGET_GENES & https://www.gsea-msigdb.org/gsea/msigdb/human/geneset/NKX2_2_TARGET_GENES), respectively, to derive brain- and pancreas-specific target lists. For PTF1A targets specific to the pancreas and brain, signature lists were sourced from Meredith et al. 2013 [28].

### Module score analysis and hypergeometric test

All module score analyses performed in this study were computed using sc.tl.score_genes(), as implemented in scanpy. Specifically, the module score was calculated by subtracting the average expression of a particular gene set from the average expression of a reference gene set. The reference set was randomly selected from the gene pool for each expression value bin. All external signature lists used for module score analysis are provided in Supplementary Table 4.

Furthermore, an over-enrichment test was conducted based on the cumulative distribution function of the hypergeometric distribution as described in https://systems.crump.ucla.edu/hypergeometric/index.php, using Scipy’s implementation in Python. The overrepresentation is quantified by the representation factor, calculated as the number of overlapping genes divided by the expected number of overlapping genes from two independent groups. A representation factor greater than 1 indicates higher overlap than expected, while a factor less than 1 indicates lower overlap. A representation factor of 1 represents the expected overlap for independent gene groups. This representation factor is calculated as follows:

- k = number of overlapping genes between the two groups (also denoted as number of successes)
- s = number of genes in group 1 (e.g., number of transition genes)
- M = number of genes in group 2 (e.g., number of DEGs)
- N = total genes expressed or tested.

The representation factor is given by *x*/expected number of genes, where the expected number of genes is defined as (*s*×*M*)/*N*.

Using GTEX bulk RNA-seq data [42] and DESeq2 analysis comparing the brain and pancreas, tissue-specific target lists for these factors were created. The shared NE and NE proliferating cell states expressed *PAX6* brain-specific targets, including *NFASC, PKD1, PRKCE,* and *STXBP1* (Figure 2, Supplementary Table 4). However, due to low overlap, the FDR was not significant.

### Classification of panNEC cell states into brain cell types/states

We performed semi-supervised classification of panNEC cell states using the Celltypist LR-based classifier, leveraging first-trimester developing brain data [15] as the reference atlas. Immune cells served as a negative control to ensure the classifier did not erroneously assign neuronal identities. When majority_voting=True, CellTypist overclusters query data via Leiden clustering (resolution=20), then assigns each subcluster the dominant predicted label from individual-cell LR scores, ensuring cluster-level transcriptional coherence. Concurrently, mode= ’best match’ directly assigns labels by selecting reference cell types with the highest LR decision scores (gene expression × model coefficients), prioritizing transcriptional similarity without probabilistic uncertainty handling. This dual approach—leveraging Leiden-driven consensus for robust subcluster identities while retaining individual-cell “best match” scores — balances population-level coherence with single-cell resolution. Subsequently, cell annotations were refined based on specific marker expression and literature-based curation of brain cell type/state markers. While amphicrine acinar cells were initially predicted as diencephalic neurons, we retained their amphicrine identity due to robust pancreatic acinar gene expression, which conflicted with a neuronal lineage.

We validated the expression of *RBFOX1* (major driving feature of brain phenotype) using the Cell*Gene platform [20]. *RBFOX1* was found to be robustly expressed across multiple neuronal and interneuronal subtypes, including both GABAergic and glutamatergic neurons, with nearly universal coverage—over 99% of cells within key neuronal clusters express *RBFOX1*, underscoring its broad and consistent neuronal specificity. Similarly, PTF1A-directed targets (especially) demonstrated strong and selective expression patterns in relevant brain cell types, further supporting their roles in neuronal development and function as identified in large-scale brain single-cell datasets (spanning around ∼18.7 million single cells). This mainly serves as an orthogonal validation of brain-specific transcriptional programs suggested from our snRNA-seq analysis.

### Integration and marker verification in adult pancreas dataset

The initial step involved integrating the panNEC snRNA-seq data with Tosti’s adult pancreas snRNA-seq dataset [19]. Stromal (normal) and immune cell types were excluded from the panNEC data. Notably, cell types originating from a common multipotent progenitor during embryonic development — including acinar and ductal cells in the exocrine compartment, as well as alpha, beta, gamma, and delta cells in the endocrine compartment were retained from the adult pancreas data and used for marker validations. Precisely, the adult pancreas dataset predominantly comprised acinar-i, acinar-s, and ductal cell types (n=42968, 31570, and 20371, respectively).

## Supporting information

panNEC_Supplementary_Tables

## Data availability

Gene expression raw count data produced by CellRanger has been deposited at GEO under accession GSE291816 (reviewer access token: qnwxqweaxvkrfip). The single-cell Scanpy object with all annotations will be made available for download via Zenodo doi: 10.5281/zenodo.15574335.

## Code availability

All analysis code required to produce the analysis and results presented in the study is available via GitHub https://github.com/HiDiHlabs/panNEC_study2025.

## Acknowledgments

We would like to thank all donors and their families. We thank Caroline Braeuning, Michaela Seeger, and Jeannine Wilde for support with single nuclei sorting, snRNAseq library preparation, and NGS. The authors acknowledge the financial support by the Federal Ministry of Education and Research of Germany in the framework of SAGE (project number 031L0265) and CNAscope (project number 01KD2443). The Pro Hominibus-Stiftung funded Bertram Wiedenmann.

