## Supplementary Information for "Aberrant brain-type neuronal programs in large-cell pancreatic neuroendocrine carcinoma"

### Supplementary information for Debnath et al. "Aberrant brain-type neuronal programs in large-cell pancreatic neuroendocrine carcinoma"

#### Extended Results

##### panNEC cell types

Using cluster-specific marker genes, we defined five main cell types: neuroendocrine (NE), amphicrine acinar, amphicrine progenitor-like, normal stroma, and immune cell types (**Figure 1c)**.

The normal stroma cell type cluster was characterized by robust and specific expression of *VIM* while lacking key neuroendocrine (NE) markers such as *SYP* (**Figure 1d & Extended Figure 1g**). This cell type also exhibited high expression of extracellular matrix (ECM) remodeling genes, including *CALD1, COL1A1, COL1A2,* and *CASC15* (**Extended Figure 1g**). Additionally, it was abundant in P1, a sample from a patient who had undergone prior chemotherapy treatments (**Extended Figure 1a, g**), while other patients showed a low abundance of this stromal cluster. This was further confirmed by immunohistochemistry (IHC, **Extended Figure 1b**).

The immune cell type cluster was characterized by prominent markers such as *PTPRC, CD247, THEMIS, ARHGAP15,* and *DOCK2*, all of which have established roles in immune biology (**Extended Figure 1g**). Both the normal stroma and immune cell types, being non-malignant, are most likely components of the panNEC tumor microenvironment (TME).

The amphicrine cell type cluster was identified based on the co-expression of the pancreatic acinar marker *GP2* [1,2] alongside neuroendocrine markers *CHGA, SYP,* and *NCAM1* (**Extended Figure 1g**). In addition to *GP2*, other key markers indicative of acinar lineage and differentiation, such as *RBPJL* [2-4, 7], *NR5A2* [4-7], *MECOM* [5-7], and *CEL,* were significantly upregulated in the amphicrine cell type compared to others (Wilcoxon rank sum test; average log2FC ≥ 0.25 and Benjamini-Hochberg adjusted p-value < 0.01) (**Extended Figure 1g**). Notably, von Figura et al. [5] highlighted *NR5A2* as a critical regulator of acinar plasticity, essential for maintaining acinar identity and reinstating acinar fate during regeneration. Similarly, *MECOM* is an oncogenic transcription factor required for acinar cell dedifferentiation [6]. The digestive enzyme trypsinogen (PRSS1) was specifically expressed in this cell type, albeit at a lower level (**Extended Figure 1g**). The amphicrine progenitor-like cell type displayed strong expression of the early pancreatic progenitor marker *PDX1* [8-10] while exhibiting negligible *GP2* expression (**Extended Figure 1g**). This cell type also demonstrated specific expression of the pancreatic progenitor marker *SOX9* [8, 10-12], along with robust expression of *FREM2, ALK,* and *RASGRF1* (**Extended Figure 1g**). *FREM2* serves as an early endodermal marker [13], typically absent in adult pancreatic cell types [3]. Moreover, *FREM2* is known to modify the extracellular matrix (ECM), creating a favorable environment for cell migration and rearrangements during embryogenesis [13].

#### Molecular landscape of panNEC shared and unique cell states

The identified cell types were further subclustered into distinct cell states (**Figure 1**) defined by robust and specific markers identified through differential gene expression analysis (average log2FC ≥ 0.25 and Bonferroni-adjusted p-value < 0.01, using multivariate logistic regression) (**Figure 1, Extended Data Figure 2, Supplementary Table 2**). Given the small number of heterogeneous samples (n = 5), no statistical test was conducted to draw definitive conclusions about the abundance or depletion of specific cell states. However, kernel density estimation (KDE) was performed to qualitatively explore compositional shifts among the patient samples (**Extended Data Figure 1e**). For instance, a noticeable trend of enrichment in the neuroendocrine proliferating cell state was observed in the P2 sample that could be associated with p53/Rb1 double mutant status (**Extended Data Figure 1a,e,f**).

The shared neuroendocrine cell state was defined by the presence of established neuroendocrine (NE) markers such as *RIMBP2* and *ADARB2*, associated with the neuronal system, along with *CACNB2, CACNA1A,* and *KCNJ3,* which are involved in voltage-gated $Ca^{2+}$ and $K^{+}$ channels, respectively [14] (**Extended Data Figure 2**). Collectively, these markers contribute to enriched pre-synaptic depolarization in the Neuroendocrine cell state (Extended Data Figure 2, Supplementary Table 2). Previously, Olofsson et al. 2024 [15] reported that the RNA editing enzyme *ADARB2* exhibits significant tissue enrichment in the brain, showing a 2-8-fold increase compared to its highest expression in non-brain tissues using Human Protein Atlas analysis. According to this study and the Human Protein Atlas, the most pronounced expression levels were observed in the spinal cord (nTPM: 31.9) and midbrain (nTPM: 20.1). Single-cell RNA expression analysis [15] in the aforementioned revealed enrichment in inhibitory neurons and oligodendrocyte precursor cells, further associating *ADARB2* with a brain-like phenotype. Of note, the calcium channel subunit encoded by CACNB2 is a membrane-associated guanylate kinase (MAGUK) protein that functions as the β2 subunit of the L-type calcium channel CACNA1C. Notably, L-type calcium channels facilitate the entry of $Ca^{2+}$ into the cytoplasm, regulating both cardiac excitability and excitation-contraction coupling (EC coupling) [16]. Albeit our data suggest substantial expression of these L-type calcium channels in LC-panNEC, their therapeutic targeting has not yet been reported for NEC and failed in pre-clinical and clinical studies in PDAC [17].

The closely related neuroendocrine proliferating cell state was characterized by *MKI67, DIAPH3, TOP2A,* and *CENPP* expression (**Extended Data Figure 2 & Supplementary Table 2**), associated with cell-cycle regulation and chromatin remodeling pathways, including the G2/M checkpoint, E2F transcriptional targets, DNA damage response (DDR), and Aurora kinase activation (**Figure 1 & Supplementary Table 2**). Importantly, the Fanconi anemia (FA) signaling pathway, involving *FANCA, FANCI,* tumor suppressor genes *BRCA1/2,* and the BRCA-interacting protein-encoding gene *BRIP1* **(Extended Data Figure 2)**, was found to be enriched within this NE proliferating cell state. Deficiencies or mutations in FA genes are known to impair DNA repair mechanisms, leading to chromosomal abnormalities and genomic instability, which are known genetic susceptibility factors for cancer [18]. Elevated expression of FA pathway genes has also been associated with chemotherapy resistance in cisplatin-resistant non-small cell lung cancer (NSCLC) [19] and ovarian cancer cell lines [20].

A shared neuroendocrine cell state, termed "NE HSP+ (hypoxic)", was identified by high expression of heat-shock protein-encoding genes, including *HSP90AA1, HSP90AB1, HSPE1,* and *HSPH1,* as well as hypoxia-induced genes such as *VEGFA, KDM3A,* and *NDRG1* (**Extended Data Figure 2**). This cell state demonstrated notable enrichment in pathways related to the heat stress response, hypoxia, mTORC signaling, glycolysis, upregulated MYC targets, and MAPK signaling (**Extended Data Figure 2 & Supplementary Table 2**). Cells exposed to environmental stressors such as hypoxia activate pathways like mTORC/PI3-AKT and initiate a heat stress response to adapt and survive [21, 22]. Overactivation of mTORC signaling promotes cell proliferation, survival, and metabolism, driving tumor growth and progression [23], a phenomenon frequently observed in pancreatic neuroendocrine tumors (panNET) [24]. The latter study also highlighted the frequent abnormal activation of mTOR, often caused by inactivating mutations in genes encoding negative regulators of the pathway or through indirect mechanisms [24]. Clinically, mTOR overexpression and its downstream targets have been associated with poor prognosis in various neuroendocrine tumors (NETs) [25, 26]. In a study by Shida et al. 2019 [27], immunohistochemical (IHC) analysis of tumor tissues revealed elevated mTOR expression in 67% of poorly differentiated neuroendocrine neoplasms (NENs), compared to 27% in well-differentiated counterparts. Similarly, Catena et al. 2011 [28] reported mTOR expression in 80%

of patients with poorly differentiated NEC, irrespective of tumor origin (including the pancreas, colon, lung, and small intestine) and proliferation rate.

Another cell state, termed "neuroendocrine stromal-like", exhibited moderate expression of *ROBO2*, low expression of neuroendocrine genes such as *KCNB2*, and stromal genes like *COL1A2* and *CALD1* **(Extended Data Figure 2)**. This cell state lacked highly specific markers to label it either as an NE or a stromal cell state, and was more of an intermediate state. In contrast, the "stroma (normal)" cell state, predominantly found in patient P1, was characterized by the expression of epithelial-mesenchymal transition (EMT) mediators, including *COL1A1, COL1A2, CDH11, FN1,* and *LAMA4* **(Extended Data Figure 2).** This cell state also showed robust expression of Hippo signaling members *YAP1* and *WWTR1* (TAZ) **(Extended Data Figure 2)**, which are known regulators of pancreatic tissue regeneration and neoplastic transformation in both pancreatic ductal adenocarcinoma (PDAC) and pancreatitis [29, 30]. Notably, *YAP1* is typically inactivated during normal endocrine specification and absent from pancreatic islets [31], but it has been established as a potential oncogene in PDAC [32]. This observation prompted further investigation into whether *YAP1* originates from the tumor subpopulation or adjacent stroma in patient P1, and was internally validated using immunohistochemistry (IHC). *CDH11* is implicated in promoting immunosuppression, extracellular matrix deposition, and desmoplastic stroma development [33]. Previous studies have also demonstrated that *CDH11* knockdown reduces cell migration in PDAC [34]. Although the transcriptional profile of the stroma (normal) cell state broadly resembles cancer-associated fibroblasts (CAFs), it lacks key CAF markers such as *FAP* and *S100A11* [35] for which we did not label it as CAF. To better understand the biological differences between the neuroendocrine stromal-like and stroma (normal) cell states, we performed a marker analysis. We detected higher expression of neural genes such as *RIMS2, PTPRN2, CACNA1A,* and *PLCG2* in the neuroendocrine stromal-like (average log2FC $\geq 0.25$, adjusted p-value $< 0.01$, Logistic Regression analysis; **Supplementary Table 2**). Thereafter, pathway enrichment analysis revealed Fc-mediated phagocytosis and PPI synapse enrichment in both stromal cell states (**Figure 1g**). As expected, stroma (normal) displayed significant enrichment in focal adhesion, NABA matrisome, ALK, and PDGF signaling **(Extended Data Figure 2 & Supplementary Table 2).** PDGF signaling activation plays a critical role in recruiting endothelial cells and promoting angiogenesis within the tumor microenvironment, thereby driving tumor growth and metastasis [36, 37]. Additionally, ECM compositional changes mediated by focal adhesion and NABA matrisome signaling, along with paracrine signaling through ALK and PDGF pathways, could influence immune cell infiltration, recruitment, and function within the tumor stroma [38]. Hence, the stroma (normal) cell state is similar to CAFs.

Lastly, the immune cell type cluster was further subdivided into lymphocytes and macrophages (**Figure 1**) and characterized by well-established marker genes (**Extended Data Figure 2**). Notably, *PLXDC2,* computationally identified as a macrophage marker, has been previously reported as a tumor microenvironment (TME)-related signature. Precisely, *PLXDC2* expression was found to be highly correlated with CD163-positive M2 macrophages [39]. A study by Tubau-Juni et al. 2020 [40] suggested that *PLXDC2* plays a significant immunomodulatory role, as its deficiency led to the production of pro-inflammatory macrophages in an ex vivo experimental system. In metastatic populations, tumor-infiltrating macrophages exhibited strong expression of CTSB, another defining marker of this cell state [41]. Lin et al. 2022 [42] demonstrated that CTSB+ macrophages suppress anti-tumor immune responses through various immune checkpoint pathways. Additionally, the DOCK8 protein, also associated with this cell state, has been implicated in regulating macrophage migration [43]. Another marker, *DOCK2*, was identified as playing a role in macrophage migration, phagocytosis, and ROS production [44].

#### *RBFOX1* is a robust marker of shared NE and NE proliferating cell states

Notably, two genes critical for neuronal development, *RBFOX1* [45] and *NKX2-2* [46], were found to be specific to the shared NE cell states and absent in the amphicrine cell states (**Extended Data Figures 1 and 2**). *RBFOX1* was identified as a differentially upregulated gene in both the NE and NE proliferating cell states (average log2FC = 1.4077 & 0.6638, respectively; Bonferroni-adjusted p-value $< 0.0001$, Logistic Regression), while it showed moderate expression in the NE stromal-like cell state (**Extended Data Figures 1 & 2**). Interestingly, *RBFOX1* is neither expressed in the embryonic nor adult endocrine pancreas under physiological conditions [47]. This observation was further validated by integrating an adult pancreas snRNA-seq dataset from Tosti et al. 2020 [3], which confirmed the absence of *RBFOX1* in native pancreatic cell types or states (**Extended Data Figure 3c**). *RBFOX1*, a neuronal splicing factor, plays a pivotal role in neuronal development and neuroplasticity by regulating alternative splicing of transcripts essential for synaptic function, axon guidance, and dendritic development [48, 49]. It is known to influence the splicing of calcium channel subunits and synaptic proteins, crucial for neuronal excitability and synaptic strength [50]. Henceforth, overexpressed *RBFOX1* levels in these two cell states not only serve as a key marker of the neuroendocrine phenotype but also suggest a mechanistic connection to calcium channel activity and observed neuroplastic gene expression.

#### Amphicrine cell states unique to sample P5

##### Early pancreatic progenitors and metaplastic acinar signatures in P5

In patient P5, a distinct amphicrine cell state, termed “Amphicrine progenitor-like,” was identified. Importantly, focal trypsin IHC staining validated the acinar-like distinction in P5, which was absent in other tumors (**Extended Data Figure 1b**).

This cell state was characterized by low but specific expression of key transcription factors involved in the differentiation of early pancreatic progenitors, including multipotent pancreatic progenitors (MPP) and tip progenitors [10, 51-53]. Notable MPP markers *PDX1* and *SOX9* were co-expressed in the amphicrine progenitor-like cell state (**Extended Data Figure 1g**). Previous lineage-tracing studies have linked *PDX1/SOX9* with MPP capable of differentiating into acinar, ductal, and endocrine lineages [54-56]. Collectively, the amphicrine progenitor-like cell state reflected an immature, stem cell-like phenotype.

#### Amphicrine progenitor-like regulated by WNT-BMP-NOTCH signaling

Repression of acinar lineage-associated genes, such as *RBPJL, GP2,* and digestive enzymes *PRSS1* and *PRSS2,* was observed in the amphicrine progenitor-like cell state, emphasizing its immature and less differentiated phenotype (**Extended Data Figure 4a & Supplementary Table 2**). Consistent with its progenitor characteristics, genes associated with pancreatoblastoma (PBL) [57] — an immature childhood tumor with multilineage features were prominently expressed in this cell state. These genes included *LEF1, LGR5, BMP4, ID1,* and *TCF7* **(Extended Data Figure 4a, Supplementary Table 2).**

Positive markers of the amphicrine progenitor-like cell state were statistically over-represented in PBL gene signatures extracted from previous studies [57] (representation factor: 18.0, hypergeometric p-value < 5.507e-08) **(Extended Data Figure 4b).** This suggested a strong involvement of the WNT-BMP-NOTCH signaling pathway, which emerged as a key feature of this cell state, reflecting dynamic interactions among these pathways. Such crosstalk is known to maintain stemness and progenitor cell properties by promoting cell survival and inhibiting differentiation in PBL [57]. Moreover, pathway enrichment analysis using the differentially upregulated markers (n = 200; average log2FC ≥ 0.25, Bonferroni-adjusted p-value < 0.01, Logistic Regression) further highlighted the critical role of WNT signaling in this cell state (**Extended Data Figure 4d & Supplementary Table 3**). Key mediators included *CTNNB1,*

*AXIN2, LGR5, DKK4, LEF1, BMP4, RNF43,* and *NOTUM* (**Extended Data Figure 4a & Supplementary Table 2, 3**). Importantly, *PROX1,* a neurogenesis driver previously inferred to be induced by WNT signaling [58, 59], was also differentially expressed in this cell state (**Extended Data Figure 3**).

Given its resemblance to early pancreatic progenitors, such as multipotent and trunk progenitors (*PDX1+/SOX9+*), the presence of PBL stemness signatures, and the absence of mature acinar markers, this cell state was annotated as "amphicrine progenitor-like." Additionally, significant upregulation of genes associated with invasiveness and cell migration, including *NOTUM, RASGRF1,* and *ALK,* was observed in this cell state **(Extended Data Figure 4a)**. These findings suggest a potentially more aggressive and invasive nature for this cell population within the tumor. Interestingly, the amphicrine progenitor-like cell state also demonstrated enrichment in ion transport processes **(Extended Data Figure 4d)**, driven by overexpressed markers such as *ADCY2, ATP1A1, SLC24A1, SLC7A11,* and *SLC22A15* (**Supplementary Table 2**). This indicates altered cellular homeostasis and metabolic dynamics, particularly in patient P5.

A notable overlap in markers was observed between the amphicrine progenitor-like cell state and the shared NE HSP+ (hypoxic) cell state (representation factor: 6.8, hypergeometric p-value < 2.833e-37). Pathway enrichment analysis of these shared markers revealed significant involvement in processes such as fluid shear stress, the presenilin PS1 pathway, the RAC1 pathway, thyroid signaling, and focal adhesion (**Extended Data Figure 4d & Supplementary Table 2**).

#### Amphicrine acinar cell states heterogeneity

Three distinct amphicrine acinar cell states— Amphicrine acinar01, Amphicrine acinar02, and Amphicrine acinar03— were identified within the amphicrine acinar cell type **(Extended Data**

**Figure 4).** These cell states were characterized by key acinar progenitor markers, including *RBPJL, GP2, NR5A2, MECOM,* and *CEL* **(Extended Data Figure 4b & Supplementary Table 2)**. *NR5A2* plays a pivotal role in regulating various stages of pancreatic development, including mechanisms relevant to pancreatic oncogenesis and the preservation of the exocrine phenotype [60]. Importantly, the absence of *NR5A2* during the final stages of acinar cell maturation is known to disrupt the expression of *Ptf1a* and *Rbpjl*, leading to incomplete differentiation of acinar cells [60].

Next, the Amphicrine acinar01 cell state exhibited elevated expression of trypsinogen genes (*PRSS1 & PRSS2*) **(Extended Data Figure 4a).** Both the Amphicrine acinar01 and Amphicrine acinar03 cell states showed significant enrichment of *PDX1*-regulated targets [61], with the highest enrichment observed in Amphicrine acinar01. Module score analysis revealed that Amphicrine acinar01 was statistically enriched in a metaplastic acinar program described by Schlesinger et al. 2020 in a K-Ras-driven mouse model of acinar-to-ductal metaplasia (ADM) [62] **(Extended Data Figure 4c)**. Differential expression analysis of Amphicrine acinar01 relative to Amphicrine acinar02 and Amphicrine acinar03 identified upregulated genes such as *CLPS, CPA1, SERPINI2, CPA2, PDGFD, CTRB1,* and *GALNT17* (average $\log_2$FC > 1.5, Benjamini-Hochberg adjusted p-value < 0.01; Wilcoxon rank-sum test) (**Extended Data Figure 4**). Moreover, the co-expression of metaplastic signatures and acinar enzyme-encoding genes (e.g., *CPA1* and *CPA2*) in the former suggests an intermediate phenotype that bridges early and late metaplastic states, a feature associated with pancreatic injury and ADM [62].

Amphicrine acinar02, in contrast, exhibited relatively higher expression of the serotonin biosynthesis gene *TPH1* [63] compared to the other two cell states (**Extended Data Figure 4**). In small bowel neuroendocrine tumors (SBNETs), TPH1 knockdown has been shown to reduce tumor size and vascularity in vivo [64]. Additionally, Amphicrine acinar02 strongly expressed extracellular matrix (ECM) remodeling genes such as *ABI3BP, FKBP9, IGFN1,* and *PROM1*, which are associated with ductal-like features (**Extended Data Figure 4**). Taken together, the amphicrine acinar cell states co-expressed markers of multiple pancreatic lineages or their progenitors, as described in transdifferentiation processes during pancreatic injury and regeneration. Such processes have been shown in mouse models to give rise to cell states with enteroendocrine or gastric differentiation profiles [65].

Moreover, a proliferating amphicrine cell state, termed Amphicrine acinar proliferating, was also identified. This state exhibited a molecular profile similar to the shared neuroendocrine proliferating cell state, characterized by markers such as *MKI67, DIAPH3, CENPP, BRCA1*, and *FANCA* alongside robust expression of acinar markers (**Extended Data Figure 4**).

Overall, stratifying panNEC into amphicrine versus neuroendocrine types may facilitate more personalized treatment approaches and improve the selection of homogeneous patient cohorts for clinical trials, thereby enhancing the likelihood of detecting treatment effects.

#### Transcriptional regulation and multi-lineage profiles of panNEC sub-states

To investigate transcriptional regulation within panNEC cell states, pySCENIC [66] and iRegulon [67] methodologies were employed (see Methods). Hierarchical clustering based on regulon activity identified two major subgroups: the amphicrine cell states in patient P5 and the shared neuroendocrine cell states (Figure 2).

Transcription factors inferred using pySCENIC in the amphicrine cell states were generally associated with physiological roles in the acinar or endocrine cells of the healthy pancreas. Tumoral transcription factor activity either exceeded normal physiological levels (e.g., *SOX13, PDX1, RUNX1, PBX1, LEF1*) or was reduced (*SOX9, HNF1B, HNF4G, MECOM, PTF1A*) **(Figure**

**2a).** The amphicrine progenitor-like cell state displayed differential regulation by transcription factors such as *SOX9, HNF1B, SOX13, ONECUT1, PDX1,* and *LEF1* (Figure 2a). Differential gene expression analysis comparing panNEC cell states to healthy adult pancreas cell types from Tosti et al. 2020 [3] (see Methods) revealed that *SOX9* and *HNF1B* were differentially upregulated in the healthy pancreas, and hence aligns with their roles in regulating pancreatic multipotent and trunk progenitors differentiation. Elevated *LEF1* activity in panNEC suggests a role in NEC initiation, potentially through maintaining stemness via transactivation of Wnt/CTNNB1-responsive genes [68, 69]. Highly proliferative amphicrine cell state shared elevated transcription factor activity for multiple *E2F* family members (**Figure 2a**). Among the amphicrine cell states, Amphicrine acinar01 and Amphicrine acinar03 demonstrated the highest activity for *PTF1A* and *MECOM* (**Figure 2a**). *PTF1A* is critical for specifying acinar cell identity [10, 51-53], while *MECOM* is essential for acinar cell dedifferentiation and pancreatic tumorigenesis [6]. Backx et al. 2021 [6] reported that *MECOM* is expressed during acinar development and reactivated during acute and chronic pancreatitis. As mentioned before, both cell states exhibited strong *HNF4A* activity, a transcription factor critical for early pancreatic progenitor gene expression and normal pancreatic development [70, 71], particularly in the context of the classical subtype of PDAC [72]. Among the amphicrine cluster, the amphicrine progenitor-like cell state demonstrated a distinct regulatory profile, characterized by lower inferred activity of acinar-related transcription factors, consistent with earlier marker and pathway analyses.

In contrast, the shared neuroendocrine cell states— NE and NE-proliferating exhibited unique transcription factor activities associated with pancreatic and brain organogenesis, including *PAX6, ISL1, SOX5, ETV1,* and *TCF3* **(Figure 2a).** The tumoral activity of these transcription factors in NE cell states was elevated compared to their physiological activity in the endocrine pancreas. Specifically, *PAX6* and *ISL1* are known to regulate pancreatic islet differentiation and the development of functional alpha and beta cells [73-78]. Additionally, *ISL1* and *PAX6* are essential for the survival and differentiation of pancreatic endocrine progenitors [75, 78]. Beyond its pancreatic roles, *PAX6* is a master regulator of neuronal development [79-81] and was upregulated in NEC compared to healthy pancreatic cell types **(Figure 2a)**. A study by Axelsson et al. 2017 [82] reported that *SOX5* knockdown leads to gene expression changes resembling type 2 diabetes (T2D) and impairs insulin secretion, calcium influx, and β-cell exocytosis. Similarly, *ETV1,* recognized as a master regulator in neuronal-derived gastrointestinal stromal tumors [83], was significantly upregulated in panNEC compared to healthy pancreatic controls.

Subsequently, to explore the transcriptional regulatory landscape of the shared panNEC cell states, iRegulon [67] was applied to analyze differentially upregulated targets (average $\log_2$FC ≥ 0.25, Bonferroni-adjusted p-value < 0.01). The analysis identified distinct transcriptional regulators for each shared cell state **(Extended Figure 3a)**. Here, the shared NE cell state was primarily regulated by *REST, SOX10*, and *RELA* **(Extended Figure 3a)**. Notably, *REST* acts as a transcriptional repressor and is recognized as a master regulator of hypoxia-induced neuroendocrine differentiation in prostate cancer cells [84]. A study by Labrecque et al. 2019 [85] on metastatic castration-resistant prostate cancer revealed that the loss of REST repressor activity drives the emergence of a tumor phenotype characterized by the expression of NE genes in the absence of androgen receptor (AR) signaling. As expected, the NE proliferating cell state was uniquely regulated by *E2F* family transcription factors and *FOXM1* **(Extended Figure 3a)**, both of which are strongly associated with tumor differentiation, proliferation, and metastasis in gastrointestinal and pancreatic neuroendocrine neoplasms (GEP NENs). On the other hand, the NE HSP+ (hypoxic) cell state exhibited significantly elevated activity of *HSF1* and *HIF1A* **(Extended Figure 3a)**. *HSF1* is known to promote cancer cell growth and survival and is associated with poor prognosis in multiple cancer types [86]. Physiologically, cross-regulation between *HIF1A* and *HSF1* has been reported, highlighting an adaptive mechanism linking the low-oxygen response to the activation of heat-shock proteins [87]. Additional regulators of the NE

HSP+ (hypoxic) cell state included *ATF5* and *NKX2-2* **(Extended Figure 3a)**. *ATF5* expression has been shown to inversely correlate with patient survival across diverse cancer types [88]. It plays a critical role in cancer cell survival and proliferation and represents a potential therapeutic target, as its inhibition selectively induces apoptosis in cancer cells while sparing normal cells [88]. Li et al. 2011 [89] reported that *ATF5* is upregulated during cellular stress, with its activity enhanced by *HSP70*-mediated stabilization. This post-translational regulation prolongs *ATF5* half-life, leading to increased transcription of downstream targets such as *BCL2*, further supporting cancer cell survival.

#### Aberrant neuronal phenotype linked to PTF1A-regulated brain targets

Previous studies using DNA methylation and mutational profiling have indicated an exocrine origin for panNEC [90], with a mutational spectrum more closely resembling that of exocrine pancreatic ductal adenocarcinoma (PDAC) than well-differentiated neuroendocrine tumors (NETs) [91, 92]. Within this context, the neuroendocrine phenotype in panNEC may arise through the acquisition of characteristics resembling either neural or islet neuroendocrine cell types. This study investigated whether any of the panNEC cell states showed tissue-specific target enrichment for key developmental transcription factors such as *PTF1A, PAX6,* and *NKX2-2,* which play distinct regulatory roles in the pancreas and brain.

*PTF1A* is indispensable for the proliferation of multipotent pancreatic progenitor cells and the specification and maintenance of acinar cells [10, 51-53]. In the brain, *PTF1A* regulates the development of inhibitory neurons and ensures balanced neuronal circuits [93-95]. However, tissue-specific target enrichment for *PTF1A* was not captured in the pySCENIC analysis, as this tool lacks tissue-specific reference databases. To address this, murine developmental signatures from the E12.5 neural tube and E17.5 pancreas, as described by Meredith et al. 2013 [96], were used to identify tissue-specific *PTF1A* targets.

Our analysis revealed that *PTF1A*-regulated pancreatic target signatures were significantly enriched in the Amphicrine acinar01 cell state and variably expressed in other amphicrine cell states (**Figure 2**). As expected, the shared NE and NE-proliferating cell states did not express pancreatic *PTF1A* target modules, as these primarily include genes encoding digestive enzymes and proteins associated with exocrine pancreatic function, such as those observed in chronic pancreatitis. Instead, the NE-proliferating cell state showed notable enrichment for PTF1A-regulated brain target signatures, including genes such as *HP1BP3, KHDRBS1, SSBP3, THRAP3, PRRC2B, ARID1A, LMNB1,* and *NCAM1* (representation factor: 5.33, hypergeometric p-value < 0.001) **(Figure 2c).**

Among these genes, *HP1BP3* promotes *WNT7B* activation through *EZH2* interaction, mediating drug resistance in glioblastoma and enhancing proliferation [97]. *KHDRBS1,* upregulated by c-Myc, contributes to cell survival and cancer stem cell vulnerability through Wnt/β-catenin signaling [98-100]. *SSBP3* impacts neuronal morphology, synaptic vesicle biogenesis, and pancreatic β-cell function [101-103], while *THRAP3* plays a role in DNA damage response and sensitivity to DNA-damaging agents [104]. Additionally, *PRRC2B* facilitates protein translation, crucial for cell cycle progression and proliferation [105]. Both *LMNB1* and *NCAM1* have been implicated in neuroendocrine phenotypes of primary prostatic tumors resembling small-cell neuroendocrine carcinoma [106].

Thereafter, we systematically assessed the expression of these *PTF1A* target genes in the Cell*Gene Brain Atlas, which encompasses ~18.7 million single cells or nuclei spanning a diversity of brain regions and cell types. Among the key targets, *KHDRBS1* displayed broad and robust expression across multiple progenitors and neuronal populations, including neural stem cells, neuroblasts, and pyramidal neurons. Notably, *KHDRBS1* was detected in significant fractions of cells in several lineages such as neural crest-derived neurons (~84.5%), retinal progenitor cells, (~65.5%) and cortical pyramidal neurons (~52.6%), with the highest proportions observed in amacrine and Cajal-Retzius cells, suggesting a foundational role in neurodevelopment and early neurogenic programs. Moreover, *HP1BP3* exhibited strong expression predominantly in glial and vascular-associated populations, most notably oligodendrocytes (~41.8%), mural cells (~45.3%), and perivascular cells (~45.8%), indicating potential involvement in glial biology and neurovascular interactions within the central nervous system. In contrast, *SSBP3* showed a more selective expression profile, being detected in specific neuronal subtypes such as L5/6 near-projecting glutamatergic neurons of the primary motor cortex (~41.9%) and L4 intratelencephalic projecting glutamatergic neurons (~41.5%). Importantly, *NCAM1*— another PTF1A brain target and established diagnostic marker for neuroendocrine neoplasms (NENs), commonly used alongside CgA/SYP, was found to be expressed at exceptionally high levels across both inhibitory and excitatory neuronal subtypes. Over 90% of cells in nearly all GABAergic and glutamatergic neuron clusters, including chandelier and cortical projection neurons, as well as glial populations such as oligodendrocytes, expressed *NCAM1*. This widespread *NCAM1* expression reinforces its significance as a developmental driver of the nervous system.

#### Extended References

63. Sinenko, S.A. *et al.* Tryptophan hydroxylase-2-mediated serotonin biosynthesis suppresses cell reprogramming into pluripotent state. *Int. J. Mol. Sci.* **24**, 4862 (2023). PMID: 36902295
64. Tow, D.H. *et al.* Inhibition of serotonin biosynthesis in neuroendocrine neoplasm suppresses tumor growth in vivo. *bioRxiv* (2023). PMID: 37066322
65. Ma, Z. *et al.* Single-cell transcriptomics reveals a conserved metaplasia program in pancreatic injury. *Gastroenterology* **162**, 604–620.e20 (2022). PMID: 34695382
66. Van de Sande, B. *et al.* A scalable SCENIC workflow for single-cell gene regulatory network analysis. *Nat. Protoc.* **15**, 2247–2276 (2020). PMID: 32561888
67. Janky, R. *et al.* iRegulon: from a gene list to a gene regulatory network using large motif and track collections. *PLoS Comput. Biol.* **10**, e1003731 (2014). PMID: 25058159
68. Zhao, B. *et al.* LEF1 enhances β-catenin transactivation through IDR-dependent liquid-liquid phase separation. *Life Sci. Alliance* **6**, e202302118 (2023). PMID: 37657935
69. Singhi, A.D. *et al.* Overexpression of lymphoid enhancer-binding factor 1 (LEF1) in solid-pseudopapillary neoplasms of the pancreas. *Mod. Pathol.* **27**, 1355–1363 (z014). PMID: 24658583
70. Ng, N.H.J. *et al.* HNF4A haploinsufficiency in MODY1 abrogates liver and pancreas differentiation from patient-derived induced pluripotent stem cells. *iScience* **16**, 192–205 (2019). PMID: 31195238
71. Bohuslavova, R. *et al.* NEUROD1 reinforces endocrine cell fate acquisition in pancreatic development. *Nat. Commun.* **14**, 5554 (2023). PMID: 37689751
72. Brunton, H. *et al.* HNF4A and GATA6 loss reveals therapeutically actionable subtypes in pancreatic cancer. *Cell Rep.* **31**, 107625 (2020). PMID: 32402285
73. Bohuslavova, R. *et al.* NEUROD1 reinforces endocrine cell fate acquisition in pancreatic development. *Nat. Commun.* **14**, 5554 (2023). PMID: 37689751
